# One Year of SARS-CoV-2: How Much Has the Virus Changed?

**DOI:** 10.1101/2020.12.16.423071

**Authors:** Santiago Vilar, Daniel G. Isom

## Abstract

SARS-CoV-2 coronavirus has caused a world-wide crisis with profound effects on both healthcare and the economy. In order to combat the COVID-19 pandemic, research groups have shared viral genome sequence data through the GISAID initiative. We collected and computationally profiled ∼223,000 full SARS-CoV-2 proteome sequences from GISAID over one year for emergent nonsynonymous mutations. Our analysis shows that SARS-CoV-2 proteins are mutating at substantially different rates, with most viral proteins exhibiting little mutational variability. As anticipated, our calculations capture previously reported mutations occurred in the first period of the pandemic, such as D614G (Spike), P323L (NSP12), and R203K/G204R (Nucleocapsid), but also identify recent mutations like A222V and L18F (Spike) and A220V (Nucleocapsid). Our comprehensive temporal and geographical analyses show two periods with different mutations in the SARS-CoV-2 proteome: December 2019 to June 2020 and July to November 2020. Some mutation rates differ also by geography; the main mutations in the second period occurred in Europe. Furthermore, our structure-based molecular analysis provides an exhaustive assessment of mutations in the context of 3D protein structure. Emerging sequence-to-structure data is beginning to reveal the site-specific mutational tolerance of SARS-CoV2 proteins as the virus continues to spread around the globe.

## 1. Introduction

The novel severe acute respiratory syndrome coronavirus 2 (SARS-CoV-2) and the resulting COVID-19 pandemic are causing a global health and economic crisis. Similar to related coronavirus, such as MERS-CoV and SARS-CoV [1, 2], SARS-CoV-2 has a 29.9 Kb positive-sense single-stranded RNA genome that encodes 29 viral components [2, 3]. Most of these components (16 total) are non-structural proteins transcribed as two large polyproteins (Orf1a and Orf1b) that are processed into individual polypeptides by viral proteases (Mpro and PLpro). The remainder of the viral proteome encodes for a variety of accessory and structural components, including the Spike (S), Envelope (E), Membrane (M), and Nucleocapsid (N) proteins.

Mutations provide the virus with mechanisms to increase the transmissibility, modify pathogenicity and evade host immunity shifting the antigenic response and causing resistance to therapeutics. SARS-CoV-2 is a RNA virus, a family with significant adaptive evolution due to high mutation rates [4]. Although the changes in coronaviruses are slower than most RNA viruses, there are some viral components in SARS-CoV-2 that already yielded relevant mutations [4-12]. In addition, there are differences in the behavior of the multiple viral components. Some proteins, such as the Spike, seem more susceptible to mutations, likely due to its pivotal role in entering the host cells and altering infectivity. The functional mean and evolutionary importance of most of the SARS-CoV-2 mutations are still being discussed. Moreover, our results indicate that, as more data becomes available, new viral mutations show up and further monitoring will be necessary to evaluate their role. Continue surveillance and knowledge of the main mutations along with their functional mean can help to reduce the healthcare impact, improve response during the pandemic and contribute to the successful development of effective vaccines and drugs that advance in the clinical process.

World-wide research groups are generating and sharing SARS-CoV-2 proteome sequence data in a rapid fashion as a global effort to combat the COVID-19 pandemic. The Global Initiative on Sharing All Influenza Data (GISAID) [13] contains more than 200,000 SARS-CoV-2 proteome sequences labelled by date and region. Sequence data is of great value to study and track the evolution and expansion of the virus. The Protein Data Bank is another crucial resource of viral protein information [14]. The 3D crystallized structure is available for multiple viral proteins, including structural proteins, such as the Spike and Nucleocapsid, the viral proteases Mpro and PLpro and some non-structural proteins like NSP12 (RNA-dependent RNA polymerase), NSP15 (Endoribonuclease) or the NSP16-NSP10 complex, among others. Combination of both resources, i.e. mapping sequence data with the available structures from the Protein Data Bank (PDB), creates an analysis tool with direct applications in the design of diagnostic tests, vaccines and drugs. Through this type of analysis, we can generate hypothesis about the crucial role of mutations in biological binding and their implication in protein function.

In this article we provide a comprehensive analysis of how much the SARS-CoV-2 virus has changed in a year using a collection of more than 223,000 proteomes. Using sequences deposited in GISAID, we quantified the mutations rates for the global proteome and the individual protein residues. Our dynamic temporal and geographical analysis identified two periods with a different mutational landscape, from December 2019 to June 2020, and from July to November 2020. The first period was critical for some previously described mutations that overtook the entire globe, such as the D614G and P323L in the Spike and NSP12 respectively [5, 8]. In the second period, additional mutations in the Spike and the Nucleocapsid were notably detectable in multiple countries in Europe. Additionally, we mapped the sequence mutations into 3D structures using the available crystallized viral proteins and further hypothesized their role and impact in protein function. These observations illuminate the correlation between sequence mutations and 3D protein structure. Our findings are in agreement with previous studies with smaller sequence sets, confirms anterior results and provides new insights about current mutations in the SARS-CoV-2 virus. Our temporal, geographical and molecular mapping has applicability in tracking viral evolution, provides surveillance support and guides the viral mechanistic comprehension crucial in the development of diagnostic tests, vaccines and drugs.

## 2. Results and Discussion

### 2.1. Components of the SARS-CoV-2 proteome are mutating at different rates

The multiple viral components behave in a different manner from a mutational perspective. In this section we analyzed the components of the SARS-CoV-2 proteome and identified the high frequency mutating viral proteins along with initial and more recent relevant residue mutations.

The SARS-CoV-2 proteome sequences, epidemiological, temporal and geographical data is available at the GISAID initiative [13]. We collected ∼223,000 full SARS-CoV-2 proteome aligned sequences from GISAID along with additional metadata from December 2019 to November 2020. For each viral protein we calculated individual residue mutation rates (MRs) and ranked residue variability to study the main viral mutations (Figure 1 and S1 of the Supplementary Material. Residue MRs are also provided in Table S1 as part of the Supplementary Material). As a measure of protein variability, we calculated the range in the residue mutation rates for each protein in the proteome. The proteins with highest range are the Spike, NSP12, NS9c and Nucleocapsid (Figure 1A). Our analysis showed that the viral components are evolving at different rates. Some proteins, like the Envelope (E) protein, have low MRs across the residue sequence while other viral components, such as the Spike (S) or the Nucleocapsid (N) showed higher degree of variability. Our results yielded some residues with higher mutation rates and confirmed some important mutations already described in the bibliography.

**Figure 1.**
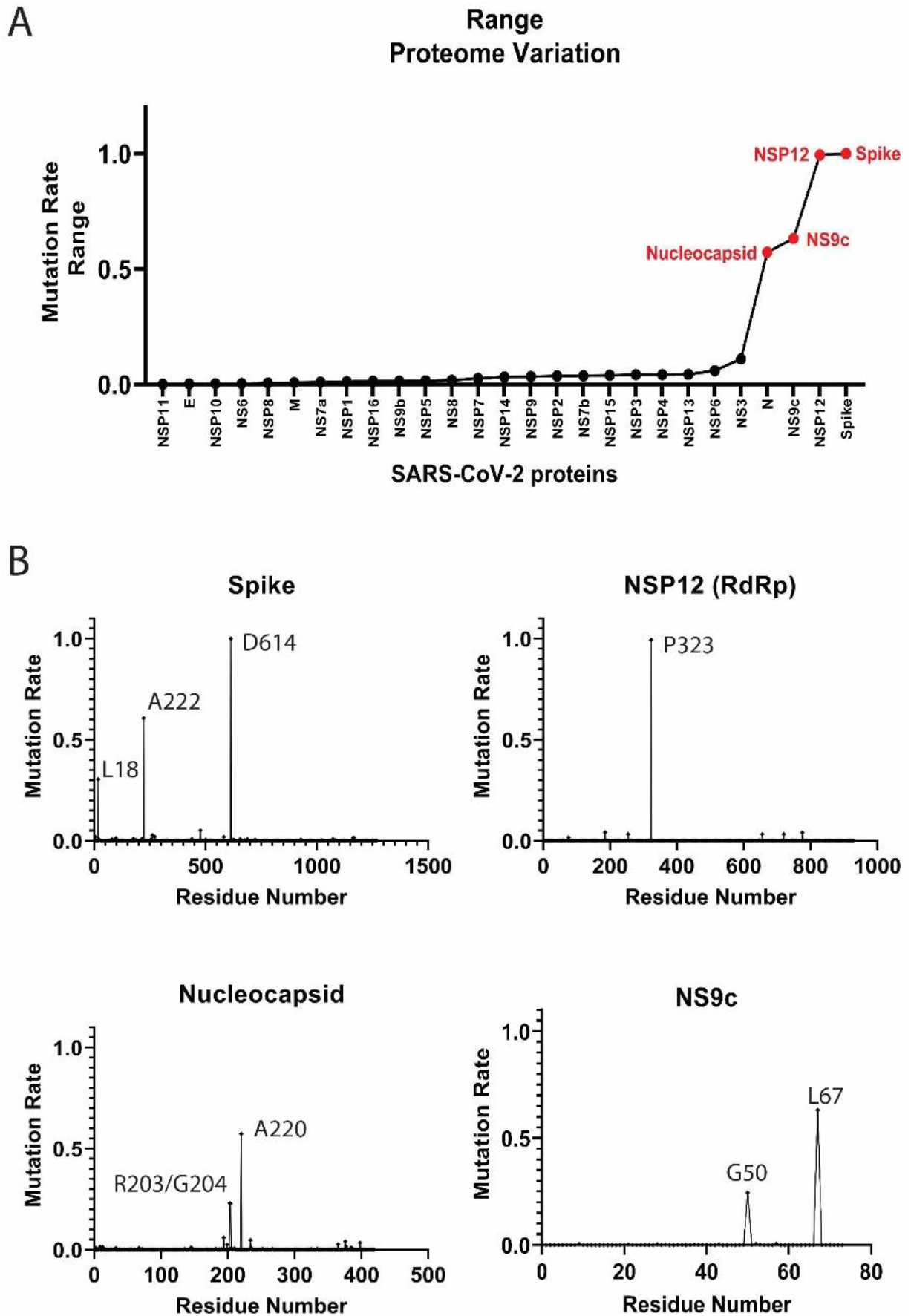
Mutation rates in the SARS-CoV-2 proteome. (A) Proteome-wide analysis of the observed mutation rate range for 27 SARS-CoV-2 proteins. Red labels correspond to proteins with a range > 0.50. (B) Select examples of high frequent mutating SARS-CoV-2 proteins. A comprehensive analysis of the mutation rates for the rest of SARS-CoV-2 proteins is available in Figure S1 of the Supplementary Material.

#### 2.1.1. High frequency mutating SARS-CoV-2 proteome components

Several SARS-CoV-2 proteins are mutating at appreciably rapid rates. While it is currently unclear if these mutations benefit the virus, their continued surveillance and the detection of new proteome variants are likely to illuminate keys aspects of viral function. As will be discussed, the identification and analysis of mutations in the Spike protein are beginning to provide such insight [5, 15]. However, the effects of mutations in other high frequency mutating viral components, such as the Nucleocapsid and NS9c proteins, is less clear. Here, we summarize the high frequency mutations that have been observed in the SARS-CoV-2 proteins.

##### Spike protein

The residue D614 of the Spike (S) protein showed a mutation rate in October-November 2020 of 0.999. The D614G mutation has already been studied in different publications [5, 9]. The Spike (S) glycoprotein mediates the entry of SARS-CoV-2 into the host cells. The D614G mutation has been associated with an increase of infectivity but not with an augment of the disease severity [5, 15]. A222V and L18F in the Spike were also mutations detected in our analysis (MRs = 0.61 and 0.31 respectively) (Figure 1B).

##### NSP12 protein

The P323L mutation in the NSP12 (RNA-dependent RNA polymerase, RdRp) protein accompanies the D614G (S) mutation in most of the analyzed sequences (MR=0.994). This dual mutation has also been reported for multiple research groups [5, 16]. As RdRp catalyzes the replication of RNA, the P323L mutation could affect the speed of the viral replication [16]. However, the P323L mutation is situated far away from the catalytic site. Other mutated residues in RdRp showed lower mutation rates, such as A185 (MR=0.04) and V776 (MR=0.04) and occupied remote positions from the pocket.

##### Nucleocapsid protein

The Nucleocapsid is another target essential in the production of viral particles, involved in RNA replication, transcription and genome assembly [17]. The Nucleocapsid also presented two consecutive residues with high mutation rates, equivalent to the mutations R203K and G204R (MRs = 0.23) (Figure 1B) [9]. Although these mutations generated lower expectation in previous literature, the residues could impact key regions for the transcription and replication of SARS-CoV-2 [18]. However, our latest data indicates that the virus is mutating back to its initial form in those residue positions. Another mutation in the Nucleocapsid, the A220V, has gained importance recently (MR=0.57). This mutation along with A222V (Spike) have been already included in a new viral variant spread in Europe during the summer 2020 [19]. The next most mutated residues in the N protein were S194 and M234 with MRs of 0.06 and 0.05, respectively.

##### NS9c accessory protein

Mutations such as G50N (MR=0.25) and L67F (MR=0.63) in the NS9c are highly correlated with residues R203/G204 and A220 from the Nucleocapsid due to possible overlapping in the reading frame.

Other viral proteins showed mutations in multiple positions, although the mutation rates are notably lower (Figure S1). Changes in residue Q57 in the NS3 (MR=0.11) [20] or residue L37 in the NSP6 (MR=0.06) should also be further monitored. More studies would be necessary to clarify their implication in the viral cycle life.

### 2.2. The temporal emergence of proteome mutations

As coronaviruses have high adaptive evolution, we expect that SARS-CoV-2 present significant temporal variations. Some factors can condition the different viral variants. Growing evidence indicates that climate and seasonal effects, including temperature, humidity, sunlight and people’s habits, can contribute to the expansion of the virus [21]. Country-specific factors, such as demography, cultural practices, social interventions, travel restrictions, quarantine policies, health care capacity and reporting and tracking mechanisms, can also alter viral expansion and variation. We analyzed the evolution of the SARS-CoV-2 virus over a year, including multiple seasons. As expected, temporal analysis yielded important variations in short periods of time. Here we provide multiple examples of temporal differences in viral protein mutation rates that exhibit a variety of behaviors.

We divided the GISAID global sequence data over several months and performed a temporal residue mutation analysis for the whole proteome and the main mutations D614G (S), A222V (S), L18F (S), P323L (NSP12), R203K (N), G204R (N), A220V (N), G50N (NS9c) and L67F (NS9c) (see Figure 2). Global analysis of the proteome temporal data showed two different periods in the mutational evolution of SARS-CoV-2. We observed two different mutational tendencies from December 2019 to June 2020 (first period) and from July to November 2020 (second period). The global results were confirmed by the detailed temporal analysis of the individual residue mutations.

**Figure 2.**
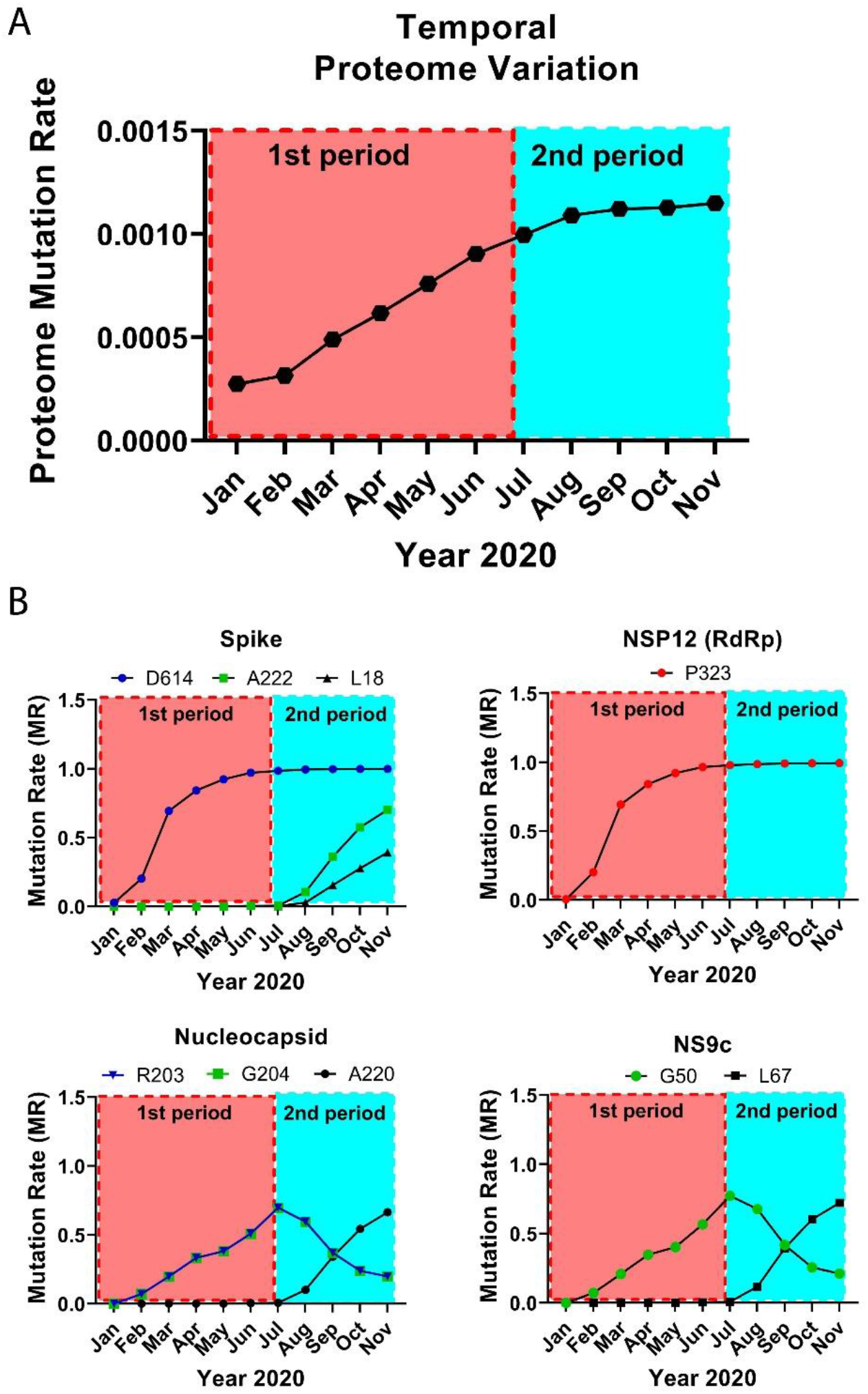
Temporal emergence of SARS-CoV-2 mutations. (A) Running temporal average of SARS-CoV-2 proteome variation relative to Dec 2019. (B) Select temporal counts of SARS-CoV-2 variation rates for the high frequency mutating residues in the Spike, NSP12, Nucleocapsid and NS9c proteins.

The virus proteome changed gradually over time reaching a maximum variation in the last analyzed month, November 2020 (global proteome MR=0.0011). However, the temporal analysis showed two periods with different slope in the mutational variation: in the first period, until June 2020, the proteome changed more abruptly and the mutation rate raised rapidly whereas the second period, from July until November 2020, showed a proteome stabilization with a slight increase in the global mutation rate (Figure 2A). Both periods are more remarkable when we analyze from a temporal perspective the main residue mutations.

When we investigated the residue mutations occurred during the first period, such as the D614G and P323L mutations, the ascend in the residue mutation rate is steeper in March with an abrupt rise in the mutation rate from 0.20 to 0.69 until a current value of ∼1 (Figure 2B). It is worth noting that the sequences collected from March to April represented more than 35% of the complete dataset. The mutation rates for residues R203/G204 of the Nucleocapsid and G50 in NS9c increased in a more gradually fashion until reaching values of ∼0.69-0.77 in July 2020. However, from that date on (second period), the MR in those residues decreased to a value of ∼0.20 in November 2020.

In our data there is a group of residues that played a more important role during the second period. The MRs of residues A222 (S), A220 (N) and L67 (NS9c) started to gradually increase in August until yielding values of ∼0.66-0.72 in November 2020. L18 (S) followed a similar pattern and reached an up-to-date MR of 0.39. These new mutations should be further monitored to establish if they play a key role in the viral life cycle.

### 2.3. World-wide geographical and temporal differences in proteome variation

As described previously, country-specific factors contribute to the viral variation and generate different patterns in the pandemic expansion. Our analysis indicated geographical differences in viral protein mutation rates and exhibited a variety of expansion behaviors. From a global perspective, we detected progressive increments in the proteome variability by country throughout the 2020. In agreement with Figure 2A, the proteome in April presents an average MR close to 0.001 in multiple countries world-wide. The proteome MR increases in most of the countries during the second period and overcomes the 0.001 threshold in September 2020 (Figure 3A).

**Figure 3.**
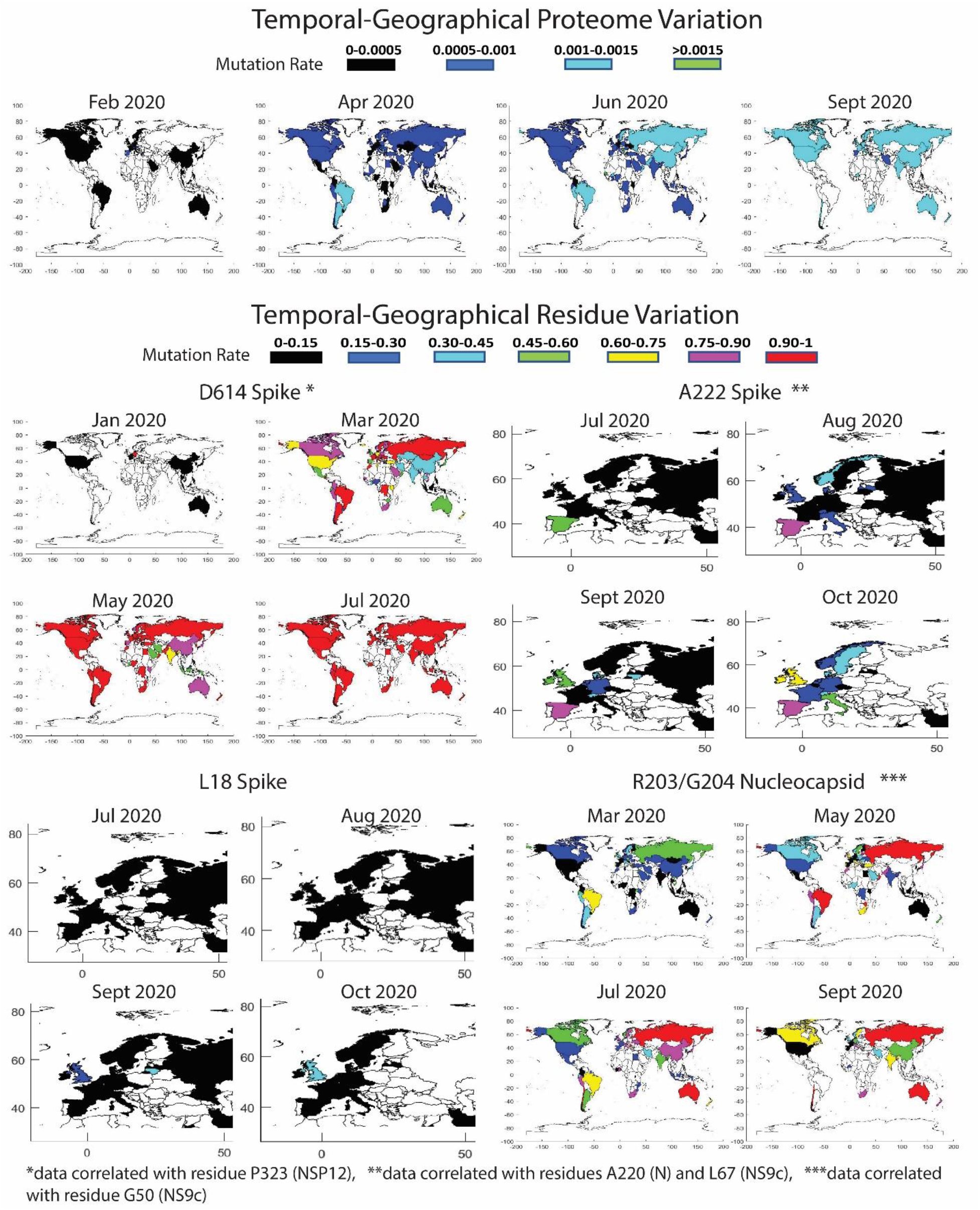
Temporal world-wide mutation rate analysis for the complete SARS-CoV-2 proteome (A) and the high frequency mutating residues (B): D614 in the Spike (correlated data for P323 in the NSP12), A222 in the Spike (correlated data for A220 in the Nucleocapsid and for L67 in the NS9c), L18 in the Spike, and R203/G204 in the Nucleocapsid (correlated data for G50 in the NS9c).

We also monitored the residue mutation rate over time in different geographic regions of the world for residues D614 (S), A222 (S), L18 (S), P323 (NSP12), R203 (N), G204 (N), A220 (N), G50 (NS9c) and L67 (NS9c) (Figure 3B). Our data showed that D614G (S) and P323L (NSP12) mutations overtook the entire globe. Mutations R203K (N), G204R (N) and G50N (NS9c) spread over the world but are less stable than the mutation D614G (S) and those residue positions were subjected to back-mutation towards the original state in multiple areas. More recent viral mutations, such as A222V (S), L18F (S), A220V (N), and L67F (NS9c) were mostly detected in Europe and should be further monitored to estimate their impact in viral evolution. A detailed description of the geographical evolution of these mutations is provided below.

#### Spike

The D614G mutation was already present in January 2020 in the sequences analyzed from Germany (MR=1, sequences=9). We detected in that period the D614G mutation in Australia and China, but the original residue was still highly conserved (MRs=0.05 and 0.01 respectively). Surprisingly, the sequences evaluated from Germany in February showed a decrease in D614 mutation (MR=0.46). As previously reported [5], half of the analyzed sequences coincided with the initial Wuhan form. However, in February other countries showed a remarkable increase in the presence of the D614G mutation, such as Saudi Arabia (MR=1), Switzerland (MR=0.97), Italy (MR=0.96), France (MR=0.81), Austria (MR=0.75), the Netherlands (MR=0.625) and Brazil (MR=0.6 but only 5 analyzed sequences). The United Kingdom and Spain showed MRs of 0.40 and 0.30. In North America, the U.S. still presented a MR for D614 of 0.06 whereas Canada showed higher evolution in this period (MR=0.33). In China, the MR was in the same range as previously reported in January (MR=0.01). There are additional countries with high D614 MRs but more representative data would be necessary to extract any conclusions (less than 5 collected sequences).

As described in Figure 2, the high increase in the incidence of the D614G mutation happened in March, where there are many countries in different areas of the globe with MR higher than 0.90, such as Estonia (MR=1), Morocco (MR=1), Argentina (MR=1), Romania (MR=1), Faroe Islands (MR=1), Italy (MR=0.99), Hungary (MR=0.98), Bosnia and Herzegovina (MR=0.96), Russia (MR=0.96), Switzerland (MR=0.95), France (MR=0.94), Croatia (MR=0.94), Brazil (MR=0.94), Denmark (MR=0.93), Luxembourg (MR=0.93), Czech Republic (MR=0.93), Costa Rica (MR=0.92), Sweden (MR=0.92) and Democratic Republic of the Congo (MR=0.91). It is worth mentioning that residue D614 showed a slower evolution in China and neighboring countries in Asia compared to the rest of the world. This situation is remarkable in April 2020 when the mutation rates for the residue were higher than 0.75 in most of the world except in some Asian countries with mutation rates between 0.3 and 0.75. After May 2020, D614 was more than 90% mutated in practically all the globe and the latest data from November 2020 shows the G614 mutated residue in practically the 100% of the sequences. Based on the difference in the temporal and geographical expansion of the mutation, we performed an enrichment analysis during April 2020 to investigate if there is an association between low D614 mutation rates and reduced mortality (number of deaths per million) in the different countries. Our goal was to investigate if the mutation could cause higher infectivity and, hence, an increase in mortality. Previous studies have shown significant correlations between the presence of D614G mutation and increased case fatality rates [22, 23]. We established different thresholds for the MRs and mortality. We detected an enrichment factor > 1.25 with associated *p*-values < 0.05 in 6 out of 12 calculations. Conversely, we only found significant results in 1 out of 12 thresholds when we looked for an association between higher MRs or presence of the D614G mutation and increased mortality. In addition, when we extended our analysis to all the residues in all the proteins (∼10,000 residues) we did not find associations between MRs and mortality. We corrected our analysis by multiple hypothesis using Bonferroni and BHFDR methods [24] and all the possible associations failed the test. More studies are necessary to prove possible associations between SARS-CoV-2 mutations and mortality.

The sequences deposited from July to November 2020 yielded new mutations in the SARS-CoV-2 (Figure 3B). According to our data, the A222V (Spike) mutation was already detected in March in Iran, in April in Turkey and in May in Mexico, Japan and Canada, among others, although the MR of the A222 residue was still low (∼0.03). However, in June 2020 the mutation is clearly detected in Spain (MR=0.42) and mildly in Senegal (MR=0.05). The mutation spread in July to Gibraltar (MR=0.2) and slightly to Norway, Belgium, Ireland and Switzerland (MRs∼0.06-0.02). The variant with A222V completely overtook Spain in August (MR=0.87) and continued its expansion to Norway (MR=0.40), Italy (MR=0.27), Latvia (MR=0.24), Switzerland (MR=0.22), the United Kingdom (MR=0.18), Denmark (MR=0.17) and other European countries (France, the Netherlands, Ireland, Germany, Sweden and Belgium). Outside Europe, the mutation was detected in China although with low rates (MR=0.05). The data in September showed that the mutation was present mainly in Spain (MR=0.86), Ireland (MR=0.51), the United Kingdom (MR=0.46), Lithuania (MR=0.44), Denmark (MR=0.35), Switzerland (MR=0.34), the Netherlands (MR=0.29), Germany (MR=0.24), Sweden (MR=0.15), France (MR=0.14), Italy (MR=0.13) and Belgium (MR=0.11). The sequences in October-November yielded an increase of the A222V mutation in multiple countries in Europe, in New Zealand (MR=0.31), Tunisia (MR=0.29) and Singapore (MR=0.1). A similar distribution pattern was found for the A220V mutation of the Nucleocapsid. Previous studies already confirmed a cluster variant with both A222V and A220V that emerged during the summer, presumably in Spain, and posteriorly spread in Europe [19].

The mutation L18F in the Spike was marginally present in different countries in March (MRs ∼0.005). The MR increased to 0.06 in United Arab Emirates in June 2020 although we only collected a sample of 17 sequences. The data showed that the mutation was residually present in multiple countries until it expanded into the United Kingdom (MR=0.07 and 6,761 analyzed sequences), China (MR=0.05, 42 sequences) and Colombia (MR=0.13 but only 8 analyzed sequences) in August 2020. We detected in September an increase in the incidence of the mutation in Lithuania (MR=0.4, 25 sequences), the United Kingdom (MR=0.23 and 14,742 sequences), Chile (MR=0.2, only 5 available sequences), Ireland (MR=0.06, 179 sequences), Sweden (MR=0.04, 54 sequences), India (MR=0.03, 119 sequences) and Singapore (MR=0.03, 33 sequences). The last data in October-November 2020 showed a MR in the United Kingdom of 0.41 (9,577sequences) and 0.14 in Norway (85 sequences) (Figure 3B). Future surveillance of the new Spike mutations is necessary to estimate the importance of the variations.

#### NSP12

The viral variant with D614G contains also the P323L mutation in the NSP12. As a result, same conclusions can be extracted for both variations. We observed a clear correlated evolution by country between residues D614 and P323 (Figure 3B).

#### Nucleocapsid

The mutations in the Nucleocapsid, located mainly in residues R203 and G204, showed different evolution patterns compared to D614G (Figure 3B). In February 2020 different European countries already displayed the R203K mutation. The residue was highly mutated in the sequences analyzed from Switzerland (MR=0.76), Austria (MR=0.75) and the Netherlands (MR=0.56), although more countries exhibited the mutation with lower mutation rates, such as Italy (MR=0.20), Germany (MR=0.16), Spain (MR=0.13), the United Kingdom (MR=0.13) and France (MR∼0.10). In this period the mutation was incipient in U.S. (MR∼0.04). In March 2020, the R203K mutation had already extended to other countries, such as Brazil, Greece, Czech Republic, Estonia, Ireland, Russia, Vietnam, among others, with a MR higher than 0.5. Nevertheless, it was in Brazil and Vietnam in April and in Lithuania, Russia, Oman and Zimbabwe in May where the R203K mutation reached the threshold of 90%. The residue evolution in the U.S. was slower but in May 2020 the mutation rate increased to 0.15. The rate increased again in June until 0.24, although the data in July showed contradictory conclusions with a lower MR of 0.19. The MR continued to decline until reaching a value of 0.13 in November. The MR decrease in the U.S. was not an isolated phenomenon and the virus after July 2020 retrieved the primitive residue in multiple countries. A similar pattern was found for residue G204 (N) with a decline in the MR in the last months in most of the countries. As described previously, the evolution of residue A220 in the Nucleocapsid is highly correlated with the data obtained for residue A222 in the Spike.

#### NS9c

Residues L67 and G50 in the NS9c showed similar expansion patterns as residues A220 and R203/G204 in the Nucleocapsid. Overlapping in the reading frame could be the cause of the highly correlated evolution detected for these residues.

### 2.4. Residue variation at 3D molecular level: mapping into crystallized proteins

The 3D analysis of the viral mutations contributes to understand the key role of specific residues, helps in the assessment of pharmacological targets and guides the design and development of novel therapeutics. We mapped the SARS-CoV-2 sequence mutations into the crystallized 3D protein structures available in the Protein Data Bank (PDB) [14]. We plotted high frequency mutations (already described throughout the manuscript) and low frequency mutations. Most of the proteins are highly conserved and the punctual mutations are not close to the main catalytic sites. Multiple viral proteins could be promising drug targets from the evolutionary perspective. Figure 4 shows the main mutations located in the 3D SARS-CoV-2 protein structures.

**Figure 4.**
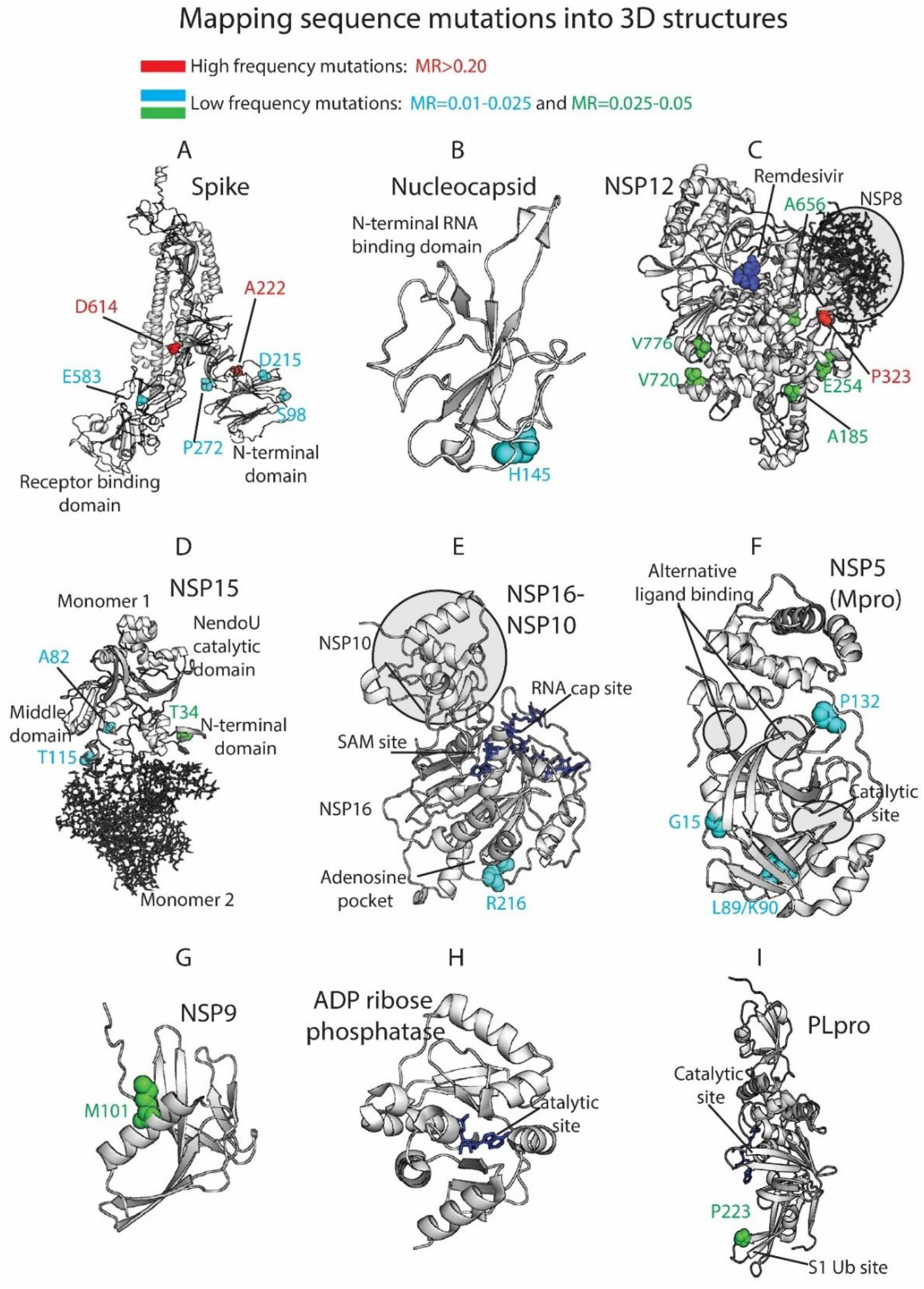
3D protein structures colored by residue mutation rates: Spike, Nucleocapsid, RdRp (NSP12), Endoribonuclease (NSP15), NSP16-NSP10 heterodimer, Mpro (NSP5), NSP9, ADP ribose phosphatase (NSP3) and Papain-like protease (PLpro, NSP3). Proteins represented in white ribbons (MRs<0.01) and color-coded residues (cyan: MRs=0.01-0.025, green: MRs=0.025-0.05, red: MRs>0.20. No residues with MR values between 0.05 and 0.20 were available in the shown crystallized structures).

The Spike (S) is a homo-trimeric transmembrane glycoprotein that mediates the viral entry into the host cells [25-27]. The Spike is the main target in the development of most of the vaccines [28] and residue variability could affect protective efficacy. The protein contains two subunits, S1 (14–685 residues) and S2 (686–1273 residues), in charge of binding to the host receptor and fusion of the host and viral membranes. The main mutation located at D614 is on the surface of each protomer (Figure 4A). The D614 established a stabilizing hydrogen bond with the residue T859 of the adjacent protomer. The mutation D614G could interrupt the mentioned hydrogen bond between both protomers, provide higher protein flexibility or even modify glycosylation at close residues, such as N616 [5]. As we have shown in our prior work [29-32], ionizable residues can be important for the pH responses of proteins, include viral components. Given the influence of pH in viral entry mechanisms [33] and the nature of D614 as an ionizable residue, the mutation could affect the pH-dependent responsiveness of the virus as it enters through the increasingly acidified endocytic pathway. One motivation for our efforts in the future will be to assess the frequency with which ionizable residues (E, C, D, H, K, R, Y) are mutated in viruses, such as SARS-CoV-2, and their role in pH-dependent endocytic entry.

The mutations A222V and L18F are far from the main D614G mutation and are located in the N-terminal domain of the S1 subunit. Alanine substitution by the bulkier valine in A222V can change inter-residue contacts and the 3D structure of the region. Both mutations D614G and A222V are located within areas defined as possible B-cell epitopes [34]. This could provide to the virus an evasive immunological advantage to avoid B-cell response. L18F is not represented in the 3D structure since the crystallized protein is missing residues M1-P26. The crystallized structure is also missing residues S477 (MR=0.05) and A262 (MR=0.03). Additional mutations with lower mutational rate are represented in Figure 4A, such as S98, D215 and P272 in the N-terminal or E583, close to the Receptor-Binding Domain (RBD), which is an essential region in the binding of the host cell receptor ACE2. Moreover, a former study in SARS-CoV associated residues 1-422 of S1 with the induction of COX-2 expression [35]. Although additional studies in SARS-CoV-2 would be necessary, mutations in this area of the S protein in the SARS-CoV-2 could be implicated in COX-2 expression and related to the inflammation response and severity of the disease.

The Nucleocapsid participates on the vital cycle of the virus in RNA assembly and release of viral particles [36]. It is an important target for pharmacological intervention, not only in the discovery of drugs but also in the development of vaccines [37]. Some of the residue mutations could interfere in the pharmacological intervention. The SARS-CoV-2 crystallized structures available at the PDB show the N2b and the RNA binding domains and do not contain key residues from the evolutionary perspective (Figure 4B). Residues R203/G204 (MR∼0.23) are not present in the crystallized structures along with A220 (MR=0.57), S194 (MR=0.06), M234 (MR=0.05), A376 (MR=0.04), A398 (MR=0.03), P365 (MR=0.03), P199 (MR=0.03) or D377 (MR=0.02), among other residues with MR greater than 0.01.

The NSP12 protein, also called RNA-dependent RNA polymerase (RdRp) is an important pharmacological target in viral intervention. Mutations in different viral RdRPs have been associated with drug resistance [38, 39]. The FDA-approved treatment for COVID-19, Remdesivir, binds the catalytic site of RdRp causing a decrease in the production of viral RNA. Our analysis showed that RdRp in SARS-CoV-2 is highly conserved as 925 residues out of 932 yielded MRs<0.005 (see Figure 4C). The residues with higher MRs, P323 (MR=0.99), A185 (MR=0.04), V776 (MR=0.04), A656 (MR=0.03), V720 (MR=0.03) and E254 (MR=0.03) are not close to the catalytic site. However, computational studies have shown that P323L and A185V mutations could have an effect in the preservation of the secondary structure of the protein that could affect protein function and drug binding [40]. Alternatively, a possible binding site was described in a hydrophobic region in close proximity to P323 [8]. RdRp forms a polymerase complex with NSP7 and NSP8 to improve RNA synthesizing activity. This complex can associate with NSP14, which is involved in replication fidelity [41]. Mutations that alter complex interactions could affect RNA replication. In fact, the mutation P323L is near the binding region between NSP12 and NSP8 and could have an impact in the polymerase complex stability (Figure 4C).

NSP15, the viral Endoribonuclease, is another possible drug target that was analyzed from the mutational point of view. The main mutated residues are T34 (MR=0.04), A82 (MR=0.02) and T115 (MR=0.02) (Figure 4D). The protein is highly conserved, and no important mutations were detected close to the catalytic site. However, some of the cited residues could collaborate in the formation of the oligomeric structure. The protein is a hexamer where the different monomers interact each other. The assembly of the hexamer is potentially sensitive to the mutations, especially in the N-terminal and middle domains [42]. T34, located in the N-terminal and T115 in the middle domain could play a role in the stabilization/destabilization of the hexamer with important implications for the Endoribonuclease functionality.

The heterodimer NSP16-NSP10 protects SARS-CoV-2 from the host immune response [43]. Additionally, disruption of NSP16 decreased the production of RNA in SARS-CoV [44]. Targeting NSP16 can facilitate immune response and decrease pathogenicity and, hence, it could be a key target in drug design. Furthermore, multiple binding sites have been described [43], including the S-adenosyl methionine (SAM) site, the RNA cap substrate cavity, and a third distant pocket unique to SARS-CoV-2 bound to adenosine. Our sequence analysis showed low mutation rates for residues in both NSP16 and NSP10. From a mutational perspective, the NSP16 pockets are highly conserved and composed of residues with MRs lower than 0.01 (Figure 4E). Residue R216 (MR=0.013) is close to the adenosine binding pocket. Important functionality of the NSP16-NSP10 complex, diversity in the binding sites and mutational stability point the heterodimer as an interesting drug target.

Another target studied by multiple research groups from the point of view of drug discovery and design is the viral main protease Mpro (NSP5) [45-48]. However, Mpro as a promising target for drug discovery against SARS-CoV-2 has risen some concerns [45]. A flexible loop constituted by residues C44-P52 can occlude the accessibility of the catalytic pocket and limit the entrance of the ligands [45]. Additionality, the plasticity of the catalytic site could make it vulnerable even to distant mutations. Our analysis identified low frequency mutations in K90 (MR=0.02), L89 (MR=0.01), G15 (MR=0.01), and P132 (MR=0.01) (Figure 4F). The cited residues are not in close proximity to either the catalytic site or two alternative binding areas described in crystallized Mpro structures (PDB_code: 5RFA, 5RGQ, 5RF0). The results showed that the main protease is a very conserved protein with high interest in drug discovery.

Other possible viral pharmacological targets yielded high degree of conservation in all the residues, such as the RNA replicase (NSP9) with role in viral RNA synthesis and viral replication [49] (all residues with MR<0.005 except M101 with MR of 0.03), the ADP ribose phosphatase, unit of the large multidomain NSP3 with possible functionality in the interference of the host immunological response [50] (MRs≤0.005 except H295 with MR of 0.01), and the PL protease, unit of the NSP3 (Figures 4G, 4H, 4I). All the PL protease residues presented MRs < 0.01 except P223 (residue P968 of the NSP3) with MR=0.03. The mutation in P223 is in the S1 ubiquitin region, one of the binding sites for ubiquitin and ubiquitin-like protein ISG15. This enzyme plays an essential role in replication and processing of viral proteins [51] but also could decrease host immunological response by collaborating in deubiquitinating and deISGylating activities [52, 53]. SARS-CoV-2-PLpro could be an excellent drug target with high residue conservation, participates in viral replication and modulates signaling in infected cells.

Low mutation rates and their important role in the virus life cycle make these proteins attractive targets for pharmacological intervention. However, according to our data, some proteins, such as the Spike or the Nucleocapsid, showed higher degree of variability in specific residues and could present future liabilities in the efficacy of vaccines and therapeutics.

## 3. Methods

### 3.1. Sequence data and residue mutation rates

GISAID database was accessed on November 30^th^ 2020 and the complete SARS-CoV-2 sequence aligned data from December 2019 was downloaded. Our database was composed of ∼223,000 sequences representing 27 viral proteins. The residue mutation rates (MRs) of the human sequences were calculated in Python [54] considering sequences with the same length, including gaps, as the original Wuhan sequences extracted in December 2019 for all the viral proteins. Residue MRs for protein *j* were computed as the ratio between the frequency in which the original residue is replaced in the protein *j* sequences and the total number of analyzed protein *j* sequences. MRs for Figure 1 were calculated comparing sequences from October to November 2020 against original sequences from China in December 2019. Protein range was calculated as the difference between the maximum and the minimum MRs.

### 3.2. Temporal analysis

Temporal fragmentation of the data was carried out extracting the sequences labeled according to the date and corresponding to consecutive months. For each period, MRs calculation was performed for each residue in the proteins and the global proteome using Python [54]. Protein variation was computed as the average of all its residue MRs. Periodical proteome variation was calculated as the average of all protein variations in each month. For clarity in the analysis, we considered two periods in the pandemic: a first period from December 2019 to June 2020 and a second period from July to November 2020.

### 3.3. Temporal/geographical analysis

Temporal and geographical fragmentation and data analysis were performed in Python and MATLAB [54, 55]. Sequence data was partitioned by date and country (∼125 world-wide countries). Multiple names representing the same country were manually inspected and unified. Residue MRs were computed as described above and plotted in world maps using MATLAB.

Association analysis between residue mutation rates and mortality in the countries were implemented with April 2020 data (period with high volume of available sequences and peak of the pandemic). We defined positive and negative cases using multiple thresholds for the mutation rates and mortality, measured as deaths per million [56]. The analysis calculated overrepresentation of countries with high/low residue mutation rates and high/low mortality. Enrichment factors with associated *p*-values were computed. All the residues were included in the analysis and *p*-values were corrected by multiple hypothesis using Bonferroni and Benjamini-Hochberg False Discovery Rate (BHFDR) methods.

### 3.4. Protein structure-based mutational analysis

Mapping sequence mutations into colored 3D crystallized proteins was performed in PyMOL [57]. The set of viral proteins with available crystallized structure in the PDB are part of the analysis. MRs for Figure 4 were computed comparing sequence data from October to November 2020 against initial SARS-CoV-2 sequences sent by December 2019.

## Supplementary Material

**Table S1**. Residue mutation rates (MRs) with values ≥0.01 for the SARS-CoV-2 proteome. Sequences from October to November 2020 were compared against the initial sequences from China in December 2019.

**Figure S1**. Residue mutation rates for the following SARS-CoV-2 proteins: NSP1, NSP2, NSP3, NSP4, NSP5 (Mpro), NSP6, NSP7, NSP8, NSP9, NSP10, NSP11, NSP13, NSP14, NSP15, NSP16, NS3, NS6, NS7a, NS7b, NS8, NS9b, Envelope (E) and Membrane (M).

## Author Contributions

Conceptualization, S.V. and D.G.I.; methodology, S.V. and D.G.I.; validation, S.V. and D.G.I.; formal analysis, S.V. and D.G.I.; writing—original draft preparation, S.V. and D.G.I.; writing—review and editing, S.V. and D.G.I.; funding acquisition, D.G.I. All authors have read and agreed to the published version of the manuscript.

## Acknowledgments

We gratefully acknowledge the GISAID Initiative along with the Originating laboratories responsible for obtaining the specimens and the Submitting laboratories where genetic sequence data were generated and shared via the GISAID Initiative.

## Abbreviations

MR: Mutation Rate
S: Spike
N: Nucleocapsid
E: Envelope
M: Membrane
RdRp: RNA-dependent RNA polymerase
PLpro: Papain-like protease
Mpro: Main protease

