## Supplementary_Material for "One Year of SARS-CoV-2: How Much Has the Virus Changed?"

**Index:**

**Table S1.** Residue mutation rates (MRs) with values ≥0.01 for the SARS-CoV-2 proteome. Sequences from October to November 2020 were compared against the initial sequences from China in December 2019.

**Figure S1.** Residue mutation rates for the following SARS-CoV-2 proteins: NSP1, NSP2, NSP3, NSP4, NSP5 (Mpro), NSP6, NSP7, NSP8, NSP9, NSP10, NSP11, NSP13, NSP14, NSP15, NSP16, NS3, NS6, NS7a, NS7b, NS8, NS9b, Envelope (E) and Membrane (M).

**Table S1.** Residue mutation rates (MRs) with values ≥0.01 for the SARS-CoV-2 proteome. Sequences from October to November 2020 were compared against the initial sequences from China in December 2019.

| Protein | Residue | Number | MR | | Protein | Residue | Number | MR |
|---|---|---|---|---|---|---|---|---|
| **Spike** | D | 614 | 1.000 | | **NSP6** | M | 143 | 0.023 |
| **NSP12** | P | 323 | 0.994 | | **NSP2** | M | 135 | 0.022 |
| **NS9c** | L | 67 | 0.632 | | **NSP6** | A | 54 | 0.021 |
| **Spike** | A | 222 | 0.606 | | **Spike** | P | 272 | 0.021 |
| **Nucleocapsid** | A | 220 | 0.572 | | **Spike** | E | 583 | 0.020 |
| **Spike** | L | 18 | 0.305 | | **Spike** | L | 5 | 0.020 |
| **NS9c** | G | 50 | 0.245 | | **NSP2** | V | 381 | 0.020 |
| **Nucleocapsid** | R | 203 | 0.230 | | **NSP15** | A | 82 | 0.017 |
| **Nucleocapsid** | G | 204 | 0.229 | | **NS7b** | H | 37 | 0.017 |
| **NS3** | Q | 57 | 0.109 | | **NS8** | A | 65 | 0.017 |
| **Nucleocapsid** | S | 194 | 0.060 | | **NSP12** | T | 76 | 0.017 |
| **NSP6** | L | 37 | 0.059 | | **NSP15** | T | 115 | 0.017 |
| **Spike** | S | 477 | 0.051 | | **Spike** | G | 1167 | 0.016 |
| **Nucleocapsid** | M | 234 | 0.047 | | **Spike** | D | 1163 | 0.016 |
| **NSP6** | L | 142 | 0.045 | | **NS3** | V | 202 | 0.016 |
| **NSP13** | H | 290 | 0.043 | | **NSP2** | L | 180 | 0.016 |
| **NSP12** | A | 185 | 0.042 | | **NSP5** | K | 90 | 0.016 |
| **NSP13** | E | 261 | 0.042 | | **Nucleocapsid** | D | 377 | 0.016 |
| **NSP4** | M | 324 | 0.042 | | **Spike** | S | 98 | 0.014 |
| **NSP3** | I | 1683 | 0.042 | | **NSP5** | G | 15 | 0.014 |
| **NSP12** | V | 776 | 0.042 | | **NS9b** | S | 6 | 0.014 |
| **NSP13** | K | 218 | 0.042 | | **Nucleocapsid** | Q | 9 | 0.014 |
| **Nucleocapsid** | A | 376 | 0.042 | | **NS8** | S | 24 | 0.014 |
| **NSP15** | T | 34 | 0.039 | | **NSP16** | R | 216 | 0.013 |
| **NSP3** | A | 1736 | 0.039 | | **NSP3** | T | 428 | 0.013 |
| **NS7b** | S | 5 | 0.037 | | **Nucleocapsid** | P | 13 | 0.013 |
| **NS3** | T | 223 | 0.036 | | **NSP3** | A | 85 | 0.013 |
| **NSP6** | K | 270 | 0.036 | | **NS3** | Q | 38 | 0.013 |
| **NSP13** | A | 598 | 0.036 | | **NSP3** | H | 295 | 0.013 |
| **NSP2** | I | 120 | 0.035 | | **NSP3** | T | 1189 | 0.013 |
| **Nucleocapsid** | A | 398 | 0.034 | | **NSP2** | A | 318 | 0.012 |
| **NSP12** | A | 656 | 0.034 | | **NSP5** | L | 89 | 0.012 |
| **NSP12** | V | 720 | 0.034 | | **Nucleocapsid** | H | 145 | 0.011 |
| **NSP3** | P | 968 | 0.034 | | **NSP3** | K | 429 | 0.011 |
| **NSP9** | M | 101 | 0.033 | | **NSP1** | D | 48 | 0.011 |
| **NSP12** | E | 254 | 0.033 | | **NSP14** | N | 129 | 0.011 |
| **NSP14** | M | 501 | 0.032 | | **NS3** | V | 163 | 0.011 |
| **NSP3** | T | 1363 | 0.029 | | **NSP6** | E | 195 | 0.011 |

| | | | | | | | | |
|---|---|---|---|---|---|---|---|---|
| **Spike** | A | 262 | 0.028 | | **Spike** | D | 215 | 0.011 |
| **NS3** | K | 75 | 0.028 | | **Spike** | L | 176 | 0.011 |
| **Nucleocapsid** | P | 365 | 0.027 | | **Nucleocapsid** | R | 385 | 0.011 |
| **NS3** | G | 172 | 0.027 | | **NSP6** | V | 149 | 0.011 |
| **NSP2** | T | 85 | 0.026 | | **NSP5** | P | 132 | 0.010 |
| **NSP6** | M | 86 | 0.026 | | **NS9b** | H | 9 | 0.010 |
| **NSP7** | M | 75 | 0.026 | | **NS3** | A | 72 | 0.010 |
| **Nucleocapsid** | P | 199 | 0.025 | | | | | |

**Figure S1.** Residue mutation rates for the following SARS-CoV-2 proteins: NSP1, NSP2, NSP3, NSP4, NSP5 (Mpro), NSP6, NSP7, NSP8, NSP9, NSP10, NSP11, NSP13, NSP14, NSP15, NSP16, NS3, NS6, NS7a, NS7b, NS8, NS9b, Envelope (E) and Membrane (M).

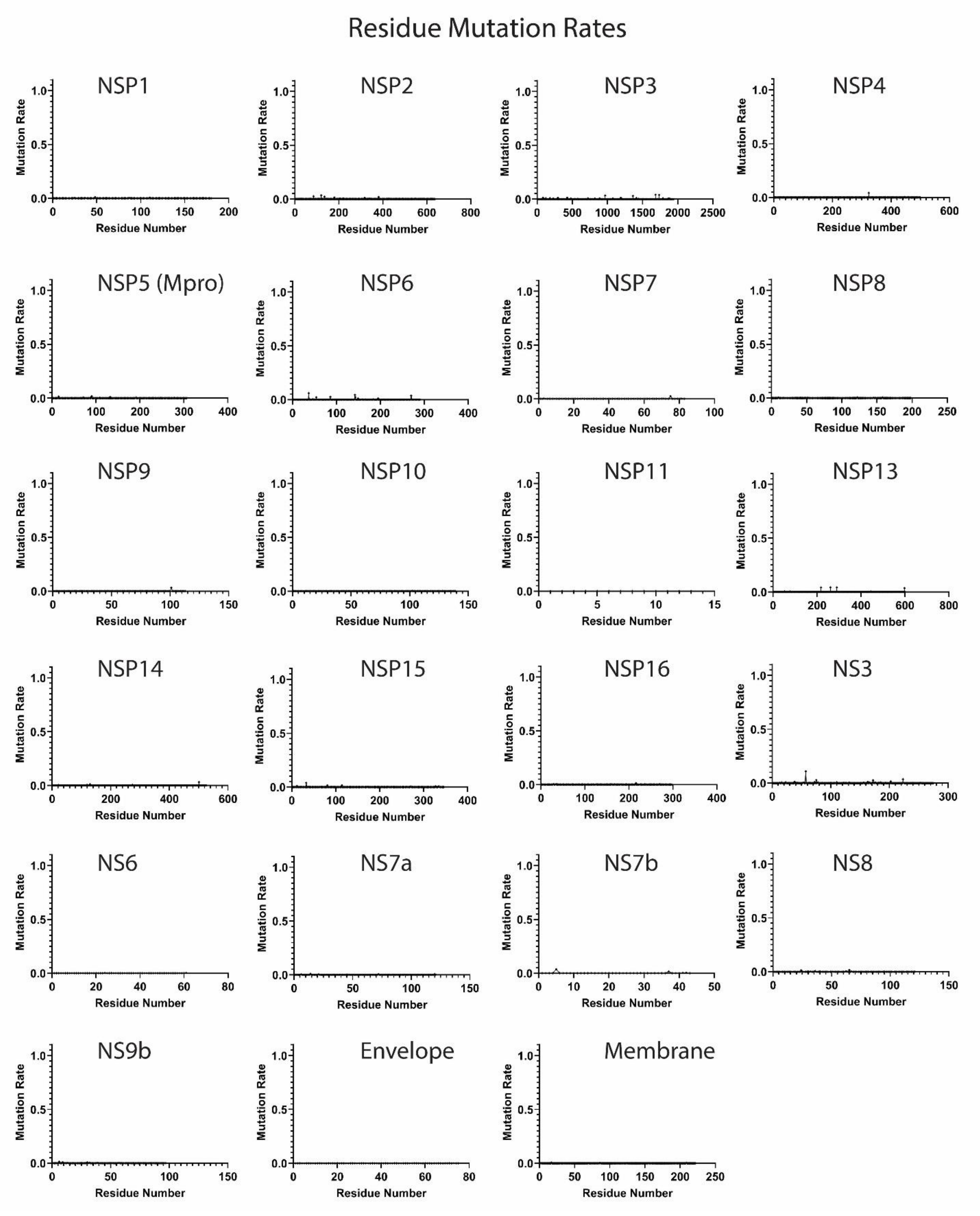
